# Integrated Techniques for Extracellular Particle Separation and Single-Particle Multiparametric Characterization to Track Cancer Biomarkers from Tissue to Biofluids

**DOI:** 10.1101/2025.01.09.632270

**Authors:** Kim Truc Nguyen, Jiangping Xia, Chaoran Chang, Mangesh Dattu Hade, Xilal Y. Rima, Ji Yeong An, Byung Hoon Min, Chiranth K. Nagaraj, Lin Wang, Tony Jun Huang, Feng Li, Yong Kim, David T.W. Wong, Eduardo Reategui

## Abstract

Gastric cancer (GC) remains a formidable global health challenge, with late-stage diagnosis and high recurrence rates resulting in poor patient outcomes. This study explores the potential of advanced technologies, namely Bessel Beam Excitation Separation Technology (BEST) and multiparametric biochip assay (MBA), to track single extracellular vesicle and particle (EVP) cargo from organs of pathology into biofluids such as plasma and saliva. Using GC as a study model, we conducted high throughput, multiparametric analyses of EVPs derived from plasma, saliva, and tissue samples. Our findings demonstrate the feasibility of these techniques in isolating and characterizing EVPs, revealing consistent EVP morphology and size across biofluids. Furthermore, differential expression patterns of the developed and validated salivary GC biomarkers, miR-140-5p and miR-301a-3p, were observed in GC patient biofluids, supporting the diagnostic relevance of EVP cargo. Notably, saliva emerged as the most promising biofluid for GC diagnosis, achieving superior Receiver Operating Characteristic (ROC) curve values compared to plasma and tissue. This study highlights the role of BEST and MBA in advancing single-EVP analysis and elucidating EVP trafficking, paving the way for future diagnostic applications of EVP cargo.

## Introduction

Gastric cancer (GC) is one of the most prevalent cancers and represents the third leading cause of cancer-related mortality worldwide, with over one million new cases diagnosed annually.^1^ In 2020, approximately 1.1 million new GC cases and 770,000 deaths were reported globally, with the highest burden observed in East Asia, particularly in China, Japan, and South Korea. In China, GC ranks as the second most common cancer, with an estimated 679,100 new cases reported in 2015 alone.^2^ Similarly, Japan and South Korea report high incidence rates attributed to prevalent Helicobacter pylori infections and dietary factors.^3^ In contrast, the United States sees lower prevalence but significant impact, with an estimated 26,500 new GC cases and 11,130 deaths in 2023, and a five-year survival rate around 35.7%.^4^

The quest for effective biomarkers for GC is imperative to enhance early detection and treatment strategies.^5^ Traditional diagnostic methods, such as endoscopy and biopsy, are invasive and impractical for large-scale screening. Additionally, the heterogeneous nature of cancer tissue poses challenges to achieving high biopsy accuracy, highlighting the need for minimal-invasive biomarkers detectable in body fluids.^6^ The minimal invasive biopsy can provide the same or even better precision, reduce the psychological impact on patients, and reduce costs.^7–9^ MicroRNAs (miRNAs) that regulate gene expression post-transcriptionally show great promise as biomarkers for various types of cancer including gastric cancer,^10–13^ due to their stability in body fluids and involvement in cancer progression.^14–16^

To overcome the limitations of traditional diagnostic methods and harness the potential of miRNAs as biomarkers, the detection of miRNAs within extracellular vesicles and particles (EVPs) has gained prominence.^17–19^ EVPs, as natural carriers of miRNAs and other biomolecules, offer enhanced stability and an abundant source of miRNAs, making them a promising candidate for non-invasive diagnostics.^20–22^ Despite the potential use of EVPs in the cancer diagnostics, current methods for isolating and characterizing miRNAs content within the EVPs are technically challenging.^23–25^

To address these challenges, this manuscript focuses on joining two advanced techniques for particle separation and *in-situ* single-particle multiparametric characterization — Bessel Beam Excitation Separation Technology (BEST) and multiparametric biochip assay (MBA). These methodologies enable precise quantification and spatial localization of miRNAs signatures within EVPs, thus providing insights into their pathological origins and systemic distribution. By applying these techniques, we aim to elucidate the trafficking patterns of EVPs from organs of pathology into biofluids, with an emphasis on plasma and saliva.

Saliva, in particular, represents an attractive yet underexplored biofluid for diagnostic applications. Its ease of collection and non-invasive nature position it as a viable alternative to plasma for routine screening. By integrating the sub-diffraction imaging capabilities of Total Internal Reflection Fluorescence Microscopy (TIRFM) with BEST’s high-sensitivity isolation resolution, this study seeks to validate the diagnostic utility of EVP cargo in both saliva and plasma. Furthermore, these technologies provide a robust platform for understanding the biophysical and biochemical properties of EVPs, paving the way for their application in precision medicine.

Through this work, we highlight the technological advancements that enable single-particle analysis of EVP cargo, linking their biomolecular signatures to specific pathological conditions. This not only advances our understanding of EVP biology but also underscores the potential of plasma- and saliva-derived EVPs as versatile diagnostic tools. Our study cohort demographics and biomarker measurement data provide a comprehensive overview of the population under study, highlighting the potential of these miRNAs detection in biofluids as noninvasive diagnostic tools for GC.

## Result

### Integration of cutting-edge technologies

In this study, we developed and implemented a detailed workflow for the isolation and high-throughput multi-parametric analysis of EVPs derived from various paired biological samples, including saliva, plasma, and cancerous tissue from the same subject (Fig. 1a). The workflow combines advanced acoustofluidic separation (AFS) technique with automated single-particle analysis systems to ensure high purity and comprehensive characterization of EVPs. To prove for the robust and fair analysis, the study was designed as a double-blind study. Three different biospecimens were collected from either healthy donors or GC patients and assigned number codes. In this study, 10 healthy donors were recruited, with 70% male, a median age of 43 years, and 20% reporting a history of smoking. Additionally, GC patients were included, comprising 50% male, a median age of 47.3 years, and 20% with a history of smoking. The codes were unblinded after all experimental data had been collected to proceed with further analysis (Table S1).

**Figure 1:**
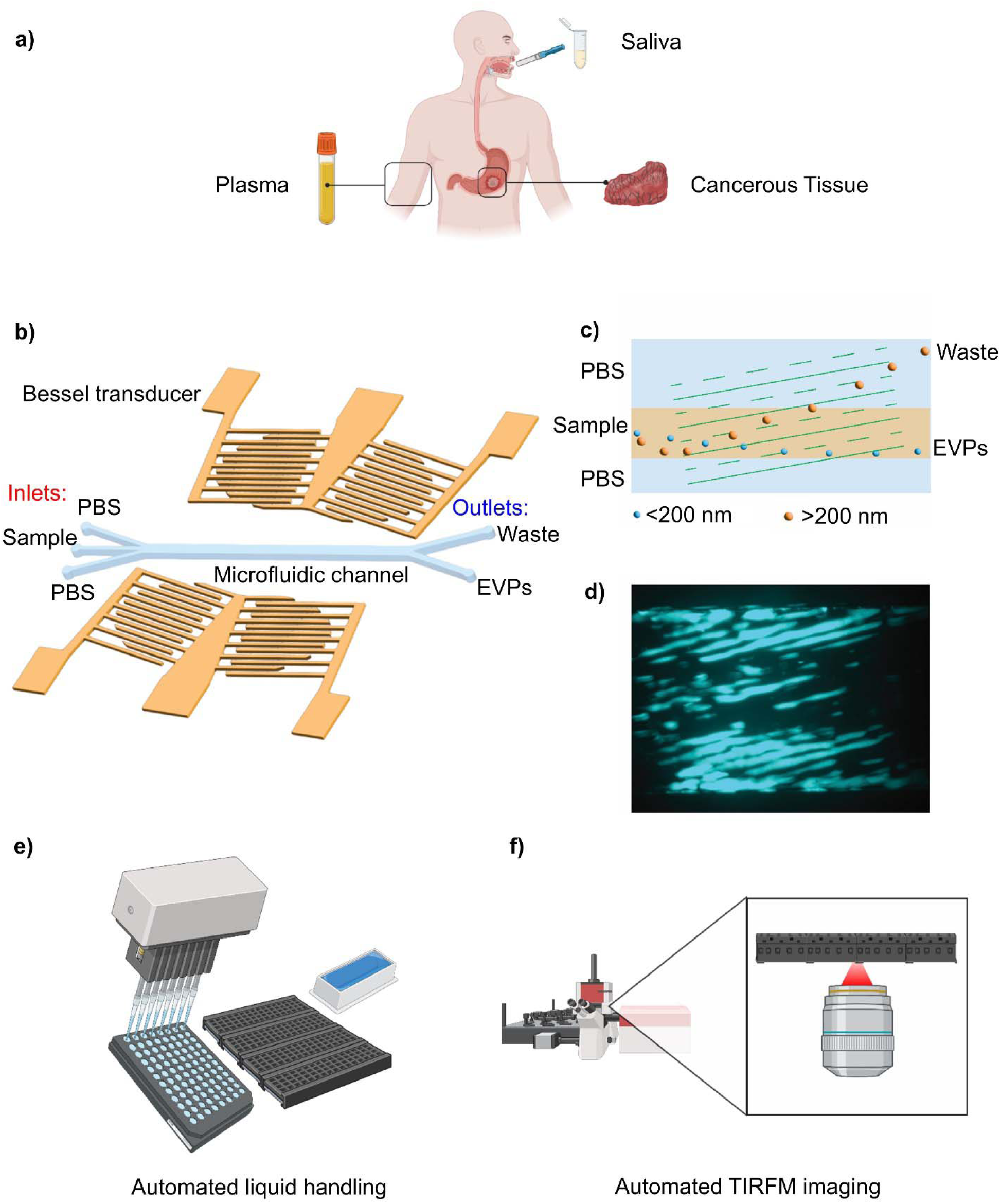
Schematic of experiment integration of techniques. (a) Orthogonal collection of three distinct biospecimens—saliva, plasma, and cancerous tissue—obtained from a single donor. (b) Schematic of the BEST chip, showing the microfluidic channel and Bessel beam transducers on a lithium niobate substrate for size-based separation. (c) Working principle of the BEST chip: larger bioparticles are directed to the waste outlet under acoustic excitation, while smaller EVPs flow to the EVPs outlet. (d) 500 nm polystyrene particles patterned by the BEST chip, demonstrating precise sub-micrometer particle manipulation. (e) Automated single-particle analysis includes an automated liquid handling machine for MBA protocol execution and (f) a total internal fluorescence microscope imaging system capable of scanning a large number of sample wells automatically.

The purification of EVPs utilized a microfluidic-based approach through acoustofluidic separation. We recently developed the acoustofluidic Bessel Beam Excitation Separation Technology (BEST), which combines acoustics and microfluidics to achieve efficient nanobioparticle isolation^36^. The BEST platform supports size-based isolation of EVPs (<200 nm) using a standing Bessel beam. This method is particularly advantageous due to its ability to process small sample volumes, achieve high throughput (30 μL/min), and preserve EVP integrity.

The BEST chip comprises two components: a pair of Bessel beam transducers and a microfluidic channel (Fig. 1b). The transducers, fabricated on a lithium niobate substrate, generate a standing Bessel beam at the channel center, creating sufficient acoustic radiation force to separate larger bioparticles (>200 nm) from smaller EVPs. The sample flow (30 μL/min) is sandwiched between two sheath flows of phosphate-buffered saline (PBS) at 45 μL/min and 15 μL/min, ensuring precise particle alignment. In the absence of acoustic activation, all bioparticles flow to the EVP outlet. When acoustics are applied, larger bioparticles experience stronger acoustic radiation forces, diverting them to the waste outlet (Fig. 1c). Demonstrating its capability, the BEST chip effectively patterns 500 nm polystyrene particles (Fig. 1d), underscoring its ability to manipulate sub-micrometer bioparticles.

Following purification, isolated EVPs underwent high-throughput analysis using automated systems designed for efficiency and precision (Fig. 1e). Automated liquid handling facilitated the preparation and processing of multiple samples simultaneously, ensuring consistency and reducing human errors. For multi-biomarkers characterization, automated TIRFM was employed. TIRFM provided high-resolution fluorescence images essential for identifying and quantifying specific EVPs markers and understanding their biological functions at the single-particle level (Fig. 1e).

This integration of microfluidic techniques for EVP purification offers significant advantages, including high efficiency, scalability, and the ability to process multiple samples in parallel. This workflow is particularly beneficial for clinical applications, where high throughput and reproducibility are essential. The combination of automated liquid handling and TIRFM imaging enhances the analytical capabilities, providing detailed insights into the molecular composition and potential diagnostic roles of EVPs.

This approach aligns with current advancements in EVP research, emphasizing the importance of innovative technologies in improving the isolation and analysis of these EVPs. The ability to isolate high-purity EVPs and analyze them at high throughput supports the exploration of EVPs biomarkers for disease diagnosis and treatment monitoring. This comprehensive workflow demonstrates the potential of combining microfluidics with automated high-throughput analysis to advance our understanding of EVPs and their roles in health and disease.

### BEST platform for size-based separation of clinical samples

BEST technique was used for the size-based separation of EVPs from various clinical samples, including plasma, saliva, and tissue. The analysis was conducted on paired sample types from a patient with GC (ID #12) and a healthy control (ID #40). The BEST technique effectively isolated the EVPs from these biofluids, allowing for detailed characterization without the interference of impurities. Transmission Electron Microscopy (TEM) was subsequently performed to observe the EVP morphologies from the different biofluids for both the gastric cancer patient and the healthy control. The TEM analysis revealed that the EVPs exhibited similar morphology (Fig. 2). Nanoparticle tracking analysis (NTA) measurement of all 60 specimens showed uniform size across three biospecimen types (d ∼ 120 – 200 nm) (Fig. 2). Additionally, there were no significant differences in EVP characteristics between the GC patient and the healthy control samples. This consistent morphology and size of EVs across different biofluids and between diseased and healthy states suggest that the BEST technique is reliable for the separation and characterization of EVPs in clinical samples. These findings contribute to our understanding of EVP’s role and consistency in various physiological and pathological conditions, highlighting their potential as biomarkers for diseases such as GC. Due to the three different sources of biospecimens, the quantity and volume of collected samples varied, resulting in differences in isolated particle concentration across sample types (Fig. S1). Therefore, particle concentrations were normalized across all samples before single-particle analysis to ensure a uniform comparison.

**Figure 2:**
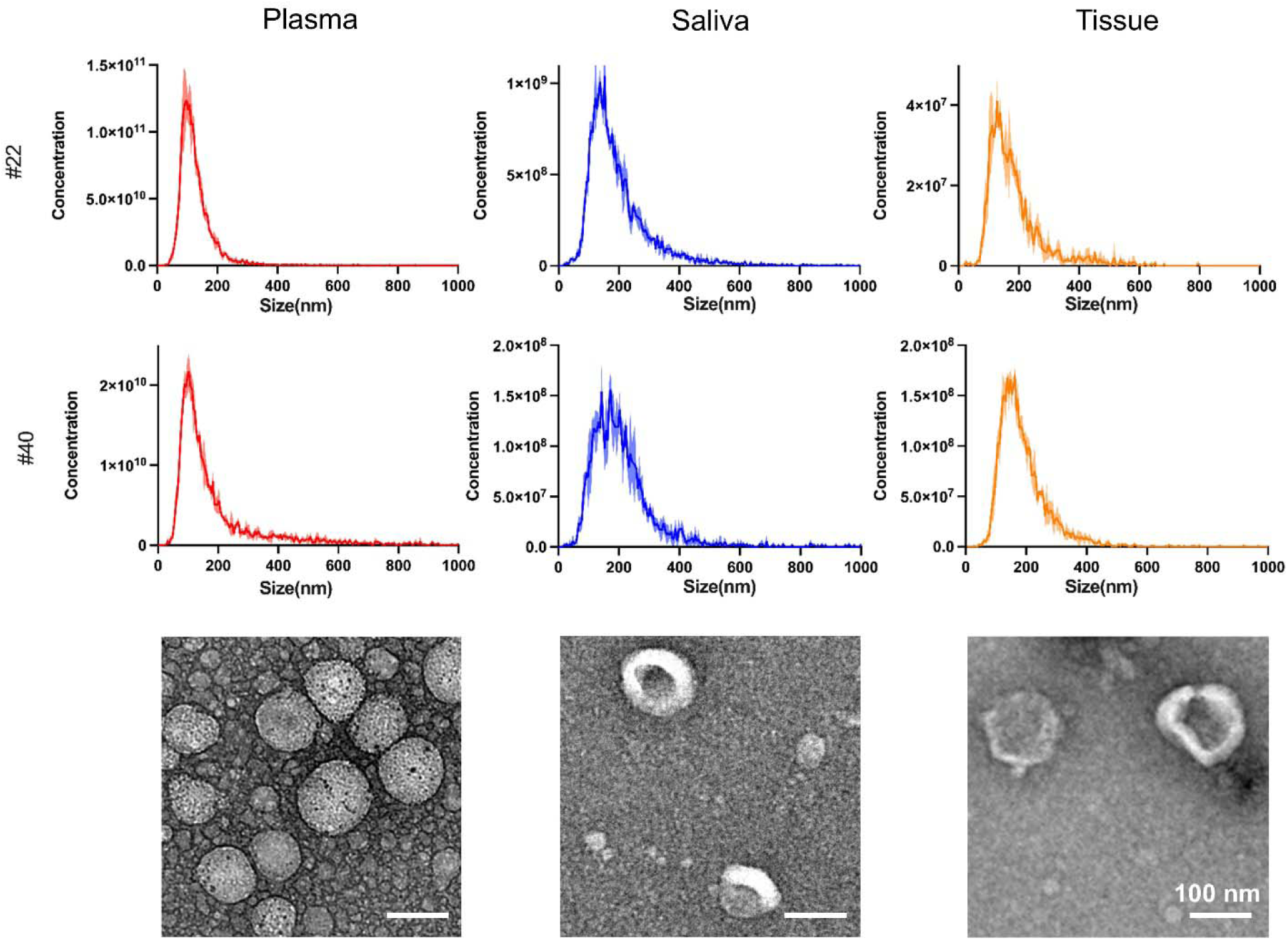
Physical characterization of isolated EVPs. Representative NTA measurement of EVPs isolated from different biospecimens, plasma, saliva, and tissue. Comparison across different sample cohort, gastric cancer and healthy control are shown. Transmission electron microscope (TEM) image showing the similar morphology of EVPs from different sample types.

### MBA for single EVPs subpopulation characterization

We employed single EVP multi-parametric characterization using a biochip functionalized with antibodies specific to EVP markers from biospecimens, including plasma, saliva, and tissue, from GC patients and healthy controls. Briefly, high-precision borosilicate glass coverslips were gold-coated and functionalized with biotin. To capture EVs, the biochip surface was coated with NeutrAvidin and biotinylated capture antibodies specific to EV markers CD63 and CD9 (Fig. 3a). The biofluid samples containing EVPs were incubated in the wells of the biochip, allowing the EV subpopulation to bind to the capture antibodies. For molecular beacon hybridization, TE buffer was used to stabilize the molecular beacons (MBs) and also permeabilize the EV membrane, facilitating hybridization with target miRNAs within the EVs^37^.

**Figure 3:**
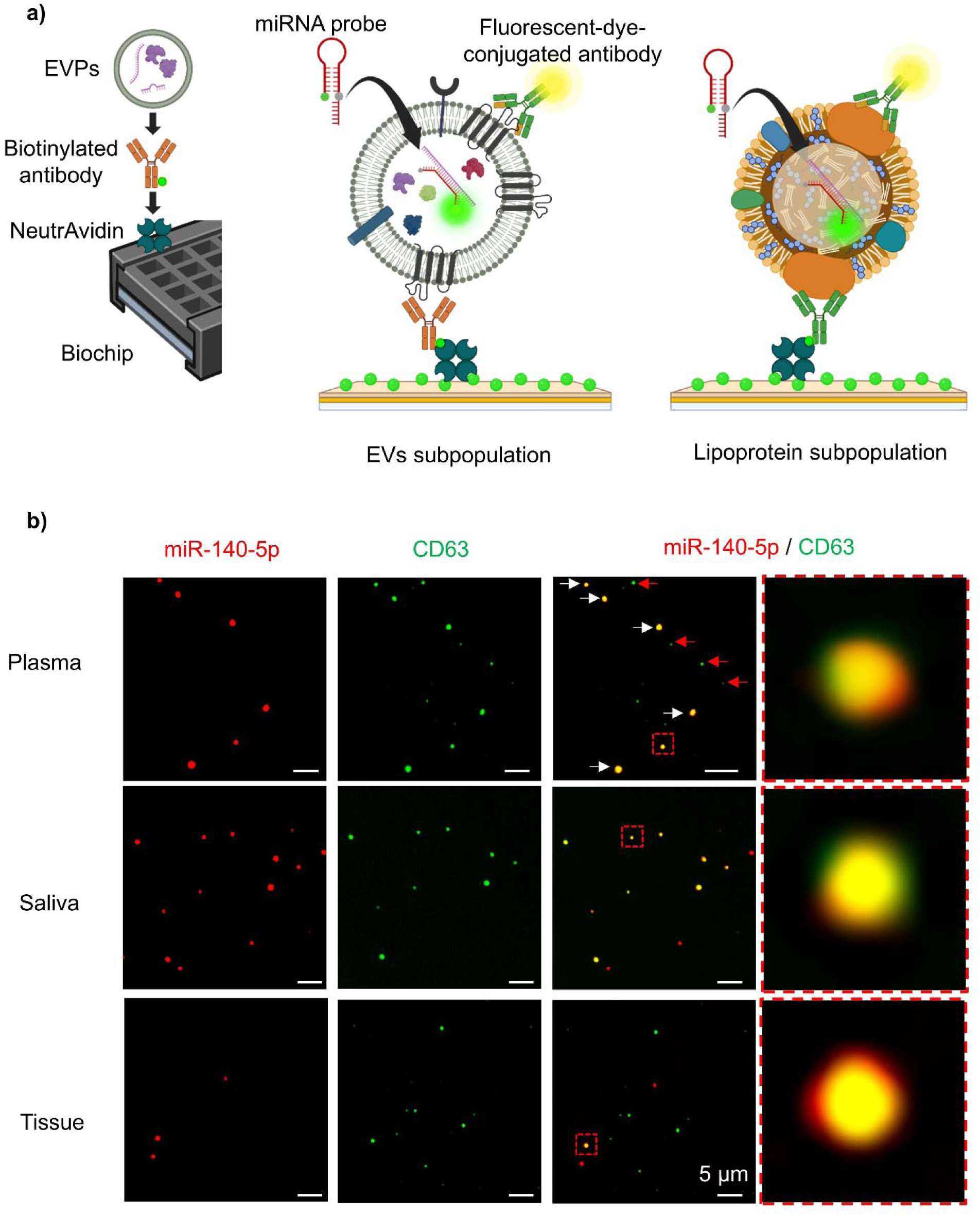
Single-particle multi-parametric analysis on biochip. (a) Schematic mechanism for detecting protein, lipoprotein and miRNA utilizing immunofluorescence and fluorescent in situ hybridization, respectively. Detection is performed on single particles with miRNA content derived from complex biospecimens, such as plasma, saliva, and tissue. (b) Representative total internal reflection fluorescence microscopy (TIRFM) images illustrating the tetraspanin CD63 common marker of EVs on single particle and miR-140-5p inside the particles. The zoom-in merged images from TIRFM illustrate the presence of anti-CD63 antibody and miR-140-5p on a single extracellular vesicle (EV) particle. The combined detection methods reveal colocalization of fluorescent signals within a specific localized region.

TIRFM was utilized to capture fluorescence images of biomarkers on the single EV. The representative TIRFM images illustrate the colocalization of miR-140-5p and CD63 within the single EV from different specimen types (Fig. 3b). Green fluorescence indicates the presence of CD63 on single particle, a common transmembrane protein marker of EV, while red fluorescence signifies the presence of miR-140-5p. Yellow signals result from the colocalization of red fluorescence of miR-140-5p and green fluorescence of CD63 within the same EV. The colocalization of miR-140-5p and CD63 was consistently observed in different biospecimens from the same subject, demonstrating the ability to effectively utilize the biochip to detect molecular markers, such as miR-140-5p, within the EV subpopulation at the single-particle level. To capture lipoprotein particles, biotinylated anti-apolipoprotein B antibody was used to functionalize on the biochip surface. Fluorescent-dye-conjugated anti-apolipoprotein B (ApoB) antibody was used as lipoprotein marker on the particles surface. Similarly, fluorescence signal of anti-ApoB antibody was colocalized with signal from miR-140-5p, showing the detection of interested miRNA in the lipoprotein particles. (Fig. S2)

These findings suggest that the detection technique is robust and that TIRFM is reliable for detailed molecular characterization of single particles that are similar in size within different subpopulations. The ability to detect and colocalize specific molecular markers within EVPs contributes to the understanding of their role in gastric cancer.

### Measuring miRNA biomarker level for gastric cancer by single EV analysis

MBA was employed for single EV molecular characterization to detect and quantify the expression of two specific GC marker miRNAs, miR-301a-3p and miR-140-5p, within EV across biospecimens, including plasma, saliva, and tissue, from GC patients (n = 10) and healthy controls (n = 10). The concentration of particles was normalized across all biospecimens to be at 1 × 10^9^ particles/mL by NTA measurement. To compare the expression level of biomarkers, total fluorescence intensity (TFI) was calculated from custom-built algorithms that were previously reported.^37,38^

Analysis revealed that TFI of EV biomarker CD63 in saliva is higher than plasma and tissue though all particles’ counts are normalized (Fig. 4a). This suggests that within the EVP populations, EVs are the majority in saliva specimens, whereas others type of particles were present in plasma and tissue. Furthermore, the expression levels of miR-301a-3p and miR-140-5p in saliva samples showed higher expression levels of these miRNAs compared to that in plasma and tissue. However, there is no clear distinction between gastric cancer patient and healthy control population when comparing the expression of single marker, either miR-301a-3p or miR-140-5p. Furthermore, the correlation of miR-301a-3p and miR-140-5p expression level separate the subpopulations between GC and healthy control (Fig. 4b). EVs derived from GC saliva express higher content of both miR-301a-3p and miR-140-5p in comparison to the healthy control. In contrast, the EVs derived from GC tissue express lower content of both miR-301a-3p and miR-140-5p. The subpopulation between GC and healthy control is not observed with EVs derived from plasmas. Receiver Operating Characteristic (ROC) curves were used to evaluate the correlation of the expression levels with GC predictions (Fig. 4c). The area under curve (AUC) values were 0.78 for saliva, 0.69 for plasma, and 0.67 for tissue, indicating that saliva had the highest predictive value for gastric cancer among the biospecimens tested.

**Figure 4:**
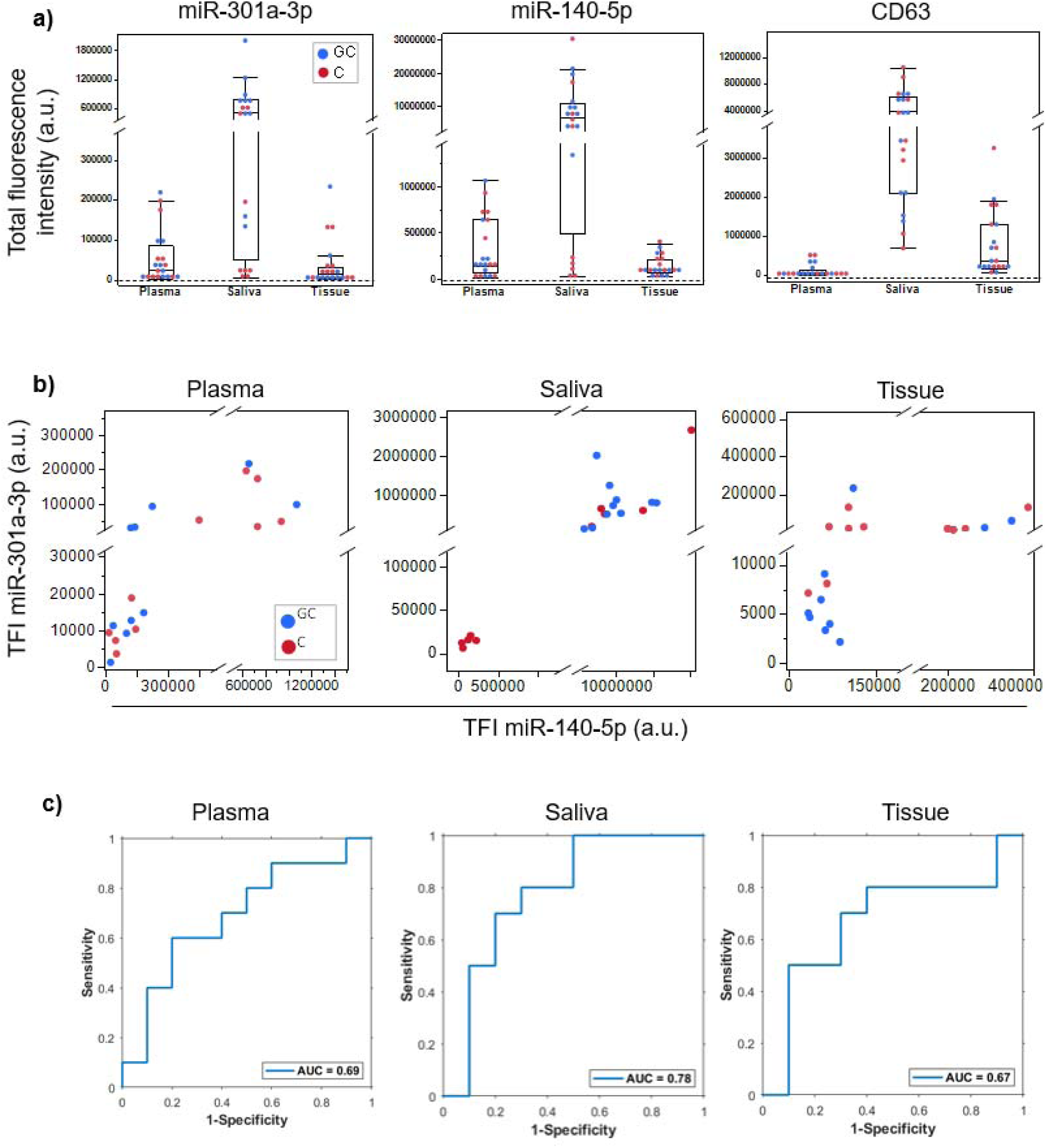
Expression level of miR-301a-3p and miR-140-5p in EVs subpopulation from three different biospecimen types. (a) Total fluorescence intensity comparison between different biospecimens using single biomarker, miR-301a-3p and miR-140-5p and EVs protein marker CD63. (b) Correlation comparison of dual miRNAs marker, miR-301a-3p and miR-140-5p. (c) Receiver operating characteristic (ROC) curves for combined detection of dual miRNAs in saliva with an enhanced area under the curve (AUC) value of 0.78.

These results demonstrate the robustness of MBA for detailed molecular characterization of single EVs and underscore the importance of miR-301a-3p and miR-140-5p as reliable biomarkers for gastric cancer. The differential expression of these miRNAs across various biofluids emphasizes the potential of EVs as non-invasive biomarkers for gastric cancer, with saliva emerging as the most promising biofluid for clinical applications. These findings lay the groundwork for the development of EV-based diagnostic tools, with the potential to transform early detection methods and enhance prognostic assessments in the treatment of gastric cancer.

### Measuring miRNA biomarker level for gastric cancer by single lipoprotein particle

The lipoprotein particles subpopulation was selectively captured and analyzed within biospecimens from gastric cancer patients and healthy controls, focusing on the expression levels of ApoB, which is highly expressed on low-density lipoproteins (LDL). LDL was hypothesized to co-isolate more effectively with EVs due to its lower molecular weight and size compared to high-density lipoproteins (HDL).

Similar biochip protocol was processed to selectively capture the LDL subpopulation using anti-ApoB antibody, and single particles within this subpopulation were analyzed. Lipoprotein particles were detected by the anti-ApoB fluorescent-dye-conjugated antibody (Fig. 3a). The TFI revealed higher levels of ApoB in plasma compared to other sample types, though the particle concentrations were normalized (Fig. 5a). This observation aligns with published findings that lipoproteins are more abundant in plasma compared to saliva and tissue^39–41^.

**Figure 5:**
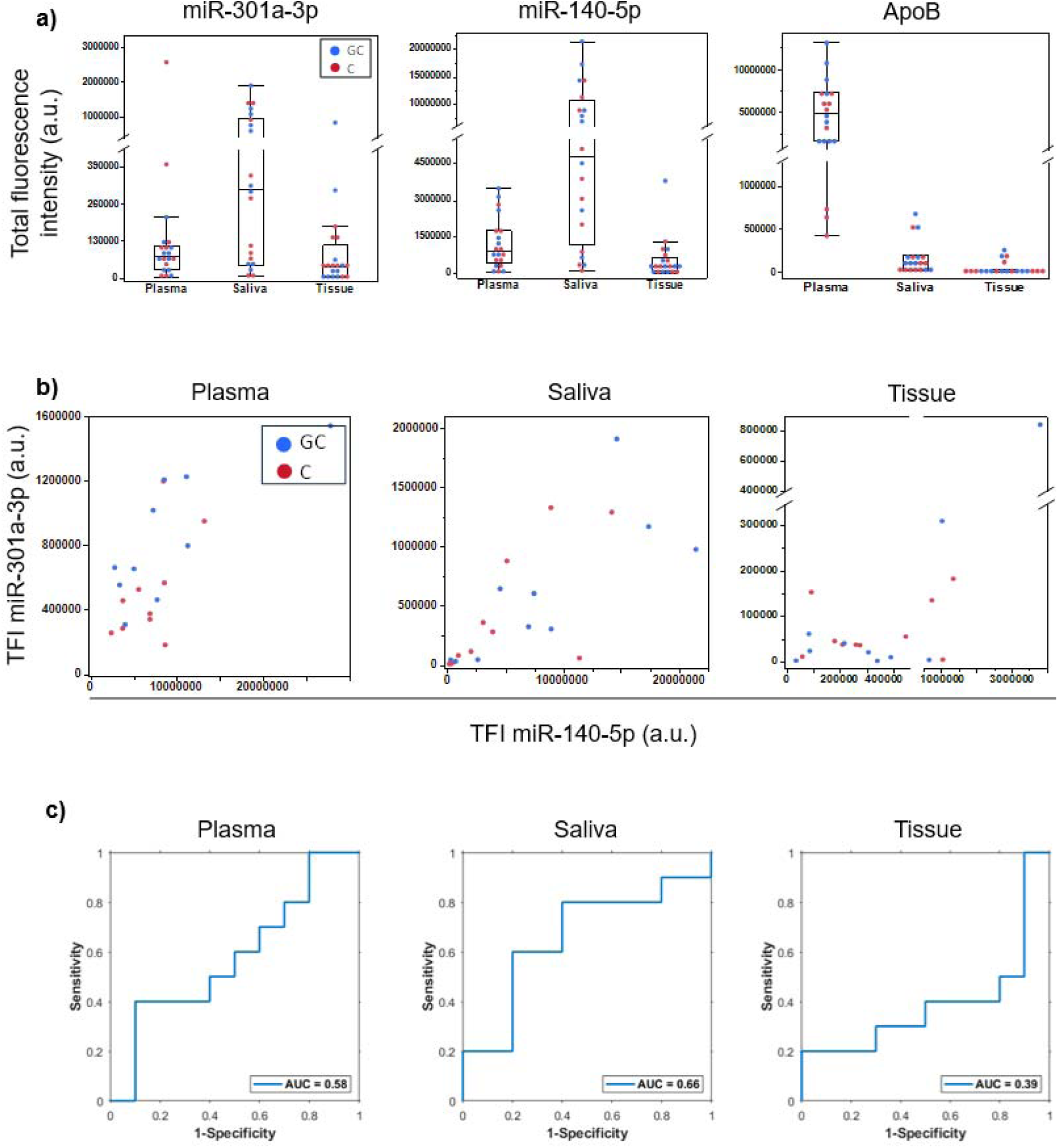
Expression level of miR-301a-3p and miR-140-5p in lipoprotein particles subpopulation from three different biospecimen types. (a) Total fluorescence intensity comparison between different biospecimens using single biomarker, miR-301a-3p and miR-140-5p and lipoprotein marker ApoB. (b) Correlation comparison of dual miRNAs marker in lipoprotein particles, miR-301a-3p and miR-140-5p. (c) Receiver operating characteristic (ROC) curves for combined detection of dual miRNAs in different biospecimens.

Despite the higher lipoprotein expressions in plasma, the expression of miR-301a-3p and miR-140-5p was higher in saliva (Fig. 5b). These findings indicate that saliva contains higher levels of these miRNAs in both EV and lipoprotein subpopulations. However, there were no distinguish between healthy control and gastric cancer samples for all type of biospecimen (Fig. 5c). ROC curves demonstrated correlations of 0.66 for saliva, 0.58 for plasma, and 0.39 for tissue, with no conclusive predictive value for GC.

These results underscore the significant content of miRNAs in lipoprotein using single-particle analysis. Though lipoprotein particles are prevalent in plasma, and they do contain significant level of miRNAs, the expression of those in lipoprotein particles do not different between healthy control and gastric cancer patient. Hence, when EVPs are analyzed using bulk technique, the different in EV subpopulation could be masked by the lipoprotein particles subpopulation. Furthermore, our method effectively detects and separates miRNAs in both EV and lipoprotein subpopulations, emphasizing its reliability for advanced analysis. This study advances the understanding of lipoprotein and EV interactions and supports the development of diagnostic tools for gastric cancer based on biofluid analysis.

## Discussion

GC presents significant diagnostic and therapeutic challenges, primarily due to late-stage detection and high recurrence rates. This study emphasizes the potential of advanced technologies, namely BEST and TIRFM, to enable precise characterization of single EVP cargo from pathological tissues and their distribution into biofluids such as plasma and saliva. By focusing on these technologies, we aimed to address the technical hurdles of isolating and analyzing EVP subpopulations. The integration of BEST and MBA demonstrated robust isolation and single-particle analysis capabilities, with consistent EVP morphology and size across biofluids. These findings affirm the reliability of the workflow and its suitability for investigating EVP cargo. The differential expression of miRNAs, specifically miR-140-5p and miR-301a-3p, across biofluids highlights the diagnostic relevance of EVPs. However, direct comparisons of diagnostic utility between tissue, plasma, and saliva should be interpreted cautiously. The differences in biomarker levels may reflect the unique characteristics of EVP cargo, which are influenced by the tissue sectioning process and biofluid-specific factors, rather than being a direct substitute for the genomic profile of heterogeneous tumor tissues.

This study was rigorously structured as a double-blind investigation to uphold the integrity and reliability of the results. Samples were collected from hospitals and encoded with unique identification numbers. Advanced analytical techniques, including BEST and biochip single-particle analysis, were applied to the samples under strict blind conditions. This approach minimized potential biases, with the blind only lifted after data collection and analysis were fully completed. Such an experimental design strengthens the validity and credibility of our findings, providing a robust basis for the observed differential expression of miR-301a-3p and miR-140-5p in GC patients.

Our findings demonstrate that miRNAs, specifically miR-301a-3p and miR-140-5p, within EVP subpopulation isolated from various biospecimens, e.g. plasma, saliva, and tissue, show differential expression patterns between GC patients and healthy controls. These results align with previous studies highlighting the role of miRNAs in cancer progression and their potential as biomarkers. miR-140-5p, known for its participation in various tumor processes^42–44^, was found to be consistently lower in GC patients, corroborating its role in inhibiting tumor proliferation, migration, and invasion through pathways such as WNT1-mediated Wnt/β-catenin signaling^29^. Conversely, miR-301a-3p, an oncogenic miRNA, was elevated in GC patients, promoting tumor progression via the PHD3/HIF-1α feedback loop^28,45^.

The utilization of advanced techniques, BEST and biochip single-particle analysis, facilitated high-purity subpopulation isolation and detailed molecular characterization of EVPs. This approach is crucial for reliable biomarker discovery, as traditional diagnostic methods like endoscopy and biopsy are invasive and impractical for large-scale screening^46^. The consistent morphology and size of EVPs across different biospecimens, observed through TEM, further validate the robustness of our isolation and analysis methods.

Saliva emerged as a promising biofluid due to its ease of collection and noninvasive nature. The expression levels of miR-301a-3p and miR-140-5p were significantly higher in saliva from GC patients compared to plasma and tissue, suggesting that saliva may be a more promising biofluid for noninvasive GC diagnosis. This finding is supported by ROC curves, which indicated higher AUC values for saliva, reinforcing its potential in clinical applications. While saliva samples demonstrated high diagnostic accuracy, this study does not assert its superiority over traditional tissue or plasma analyses. Instead, we emphasize that the focus is on demonstrating the feasibility of using EV cargo from saliva and plasma for biomarker discovery, supported by the advanced resolution and sensitivity of TIRFM and BEST. The observed diagnostic performance of saliva-derived EVs should be further validated with larger cohorts and comprehensive analyses.

Additionally, our study aligns with recent advancements in EVP research, which emphasize the importance of innovative technologies in improving EVP isolation and analysis^47^. Combining microfluidics with automated high-throughput analysis has shown significant advantages, including high efficiency, scalability, and the ability to process multiple samples in parallel. These advancements support the exploration of EVP cargo for disease diagnosis and treatment monitoring.

Moreover, our approach to capturing and analyzing the ApoB lipoprotein subpopulation within biofluids further enhances the understanding of lipoprotein and EVP interactions. The higher levels of ApoB detected in plasma compared to other biofluids align with previous reports, highlighting the distinct molecular compositions of different biospecimens^41,48–52^. This observation underscores the potential of saliva as a biofluid for miRNAs biomarker detection for GC.

This study’s incorporation of advanced isolation techniques and rigorous validation methods lays a strong foundation for the development of noninvasive diagnostic tools for GC. The differential expression of miR-301a-3p and miR-140-5p across various biospecimens, especially saliva, highlights their potential as reliable biomarkers. These findings contribute to the growing body of evidence supporting the use of EVPs in cancer diagnostics and underscore the need for further research to translate these discoveries into clinical practice. By enhancing early detection and patient management, this research holds promise for improving clinical outcomes for GC patients. In conclusion, this work establishes a technological foundation for single-particle analysis and highlights the utility of BEST and MBA in characterizing EVP cargo. By leveraging GC as a study model, we provide insights into EVP trafficking and molecular profiling that can inform future diagnostic and therapeutic strategies.

## Conclusion

This study provides compelling evidence for the diagnostic utility of EVP biomarkers, specifically miR-301a-3p and miR-140-5p, in the early detection of GC. Utilizing sophisticated isolation methodologies such as BEST and MBA for single-particle analysis, we performed an extensive multi-parametric characterization of EVPs from plasma, saliva, and tissue samples of GC patients and healthy controls. Our findings demonstrate that the expression levels of miR-301a-3p and miR-140-5p are significantly altered in GC patients, with saliva samples providing the highest diagnostic accuracy. The superior ROC curve values for saliva over plasma and tissue samples highlight saliva as the most promising biofluid for GC noninvasive diagnosis. The results highlight the promise of EV-derived miRNAs as effective biomarkers for GC. By integrating cutting-edge isolation techniques and detailed molecular analysis, this study advances our understanding of EVPs in cancer diagnostics and sets the stage for the development of novel, noninvasive diagnostic tools. This approach could greatly enhance the early identification of GC, optimize patient treatment strategies, and improve overall prognoses. Further research is warranted to validate these findings and facilitate their translation into clinical practice, ultimately aiming to enhance the diagnostic landscape for GC.

## Materials and Methods

### Saliva, plasma, and tissue collection and processing

Patient consent, recruitment, gastroscopy, and diagnostic outcome measurements were conducted at Samsung Medical Center (SMC) in Seoul, Korea. The study received IRB approval from both UCLA and Samsung Medical Center (UCLA IRB#10-000505, SMC IRB# 2008-01-028) and all experiments were performed in accordance with relevant guidelines and regulations. All patients scheduled for an endoscopic examination, endoscopic resection, or surgery at SMC of age >21 yrs. were recruited. The informed consent was administered by the attending nurse or by the respective clinical research coordinator. Informed consent form (ICF) was signed for by the recruited subject. Saliva and plasma samples were prospectively collected from 10 patients diagnosed with gastric cancer (GC) and 10 healthy donors prior to endoscopic examinations. Saliva collection was performed as described previously.^53^ About 1 mL of unstimulated whole saliva was expelled into a 50cc conical tube placed on ice. Processing occurred within 30 minutes, involving centrifugation at 2,600 ×*g* for 15 minutes at 4°C. The resulting supernatant was transferred to a 2 mL cryotube. 1 μL of Superase-In (Ambion) was added to the samples, followed by gentle inversion for thorough mixing. The cryotube was then frozen with dry ice and stored at −80°C. Whole blood sample was collected in K_2_EDTA tubes. Following the vendor’s instructions, the whole blood underwent centrifugation at 5,000 ×*g* for 15 minutes, and the resulting plasma was separated using a plasma extractor and immediately frozen for storage at −80°C. Tissue biopsy was extracted during the endoscopic procedure and snap-frozen in liquid nitrogen before storing at - 80°C.

### EVP enrichment from tissue

EVP enrichment was carried out with minimal cell lysis and disruption of exosome integrity through gentle tissue manipulation. The extraction process followed a previously described method.^54^ Briefly, the frozen human tissue (−80°C) was sliced lengthwise on ice into 0.5–1 cm long and 2–3 mm wide sections using a razor blade. Tissue sections were weighed and transferred partially frozen into a 5 mL tube containing prewarmed DMEM medium (800 µL/mg of tissue) supplemented with 75 U/mL collagenase type 3 (Worthington Biochemical) in Hibernate-E Medium (ThermoFisher). The mixture was incubated at 37°C in a shaking water bath for 20 minutes, with gentle mixing by pipetting every 5 minutes. Following incubation, the tissue was chilled on ice, treated with Complete Protease Inhibitor (SigmaAldrich) and PhosSTOP (SigmaAldrich) to a final concentration of 1x. The dissociated tissue was spun at 300 × g for 5 min at 4°C. The resulting supernatant was collected and stored at −80°C for downstream applications.

### BEST experiment

The samples were centrifuged at 800 rpm for 15 minutes to remove microscale cells and cell debris, ensuring the microfluidic channels remained unblocked during experiments. The chip design follows a previously established framework. During operation, the BEST chip was maintained at 4°C using a cooling plate. The Bessel transducers were connected to a function generator (E4422B, Agilent, USA) and a radio frequency amplifier (25A250A, Amplifier Research, USA) to generate the standing Bessel beam for EVP purification. Both transducers were driven by the same function generator at a frequency of 39.45 MHz.

### NTA

Nanoparticle Tracking Analyzer (Zetaview PMX120, Particle Matrix, Germany) was used to characterize the quality and size distribution of the isolated EVs. Isolated samples were diluted in PBS to a final volume of 5 mL, taking care to ensure that the number of particles per position detected by the NTA in each experiment ranged from 100 to 200. Sample dilutions for the 60 samples ranged from 100x to 10,000x. Three individual measurements with eleven imaging position per measurement were carried out for each sample, and the final particle diameter and concentration histograms were averaged from the three measurements.

### TEM

Negative stain transmission electron microscopy (TEM) was employed to characterize EVPs. Two 20 µL droplets of water for injection (WFI) and two 20 µL droplets of negative stain (UranyLess EM stain, Electron Microscopy Sciences) were placed on a strip of parafilm. The TEM grid underwent plasma treatment for 1 minute, after which 10 µL of the sample solution was carefully applied to the treated grid surface. The solution was allowed to incubate on the grid for 1 minute before gently blotting with filter paper to remove excess liquid. The sample grids were then rinsed by submersion in the first WFI droplet, followed by blotting, and repeated with the second WFI droplet. The grids were subsequently stained by submersion in the first droplet of negative stain, blotted, and then immersed in the second negative stain droplet. After an incubation period of approximately 22 seconds, the excess stain was gently wicked away with filter paper. For thorough drying, the stained grids were stored in a grid box overnight. TEM imaging was then performed using a Tecnai TF-20 microscope operated at 200 kV.

### The biochip fabrication

High-precision borosilicate glass coverslips (24 × 75 × 0.15 mm, D 263® M; Schott AG, Mainz, Germany) were initially cleaned in an ultrasonic bath, using ethanol followed by deionized (DI) water, for 5 minutes each. After repeating the cleaning procedure, the coverslips were dried with nitrogen gas and further treated in a UV-ozone cleaner (Jelight, Irvine, CA) for 15 minutes. A 2-nm titanium film was then deposited onto the cleaned coverslips via electron beam evaporation (DV-502A; Denton Vacuum, Moorestown, NJ) to promote gold adhesion. Using the same method, a 10-nm gold layer was deposited onto the titanium. To functionalize the gold surface with biotin, the gold-coated coverslips were immersed in a thiolated solution and incubated overnight at room temperature in the dark follow the reported method^55^. Excess thiolated molecules were then rinsed off with ethanol, after which the biotin-functionalized, gold-coated coverslips were dried with nitrogen gas and secured to a 64-well ProPlate® microarray system (Sigma-Aldrich, St. Louis, MO).

### Antibody functionalization of the biochip surface

A working volume of 20 μL was used per well and kept consistent for each solution added. All incubation steps were performed on a shaker to ensure uniform coating across the well surface. Prior to antibody functionalization, each well was washed with deionized (DI) water. A 50 μg/mL solution of NeutrAvidin (NA; Thermo Fisher Scientific), prepared in phosphate-buffered saline (PBS; Thermo Fisher Scientific), was then added to each well and incubated at room temperature for 1 hour to bind to the biotin motifs functionalized on the gold biochip surface. Excess NA was removed by rinsing with PBS and repeated three times. Biotinylated capture antibodies (Table 2) were diluted to a concentration of 10 μg/mL in a 1% (w/v) bovine serum albumin (BSA; Sigma-Aldrich) solution in PBS. This biotinylated antibody cocktail was introduced to each well and incubated at room temperature for 1 hour. Excess proteins were then removed by rinsing with PBS for three times.

### Molecular beacon hybridization to membrane-enveloped RNA

The molecular beacons (Table 3) were prepared at a concentration of 5 μM in 12.5× Tris EDTA (TE) buffer (Sigma-Aldrich), diluted with deionized (DI) water to stabilize the beacons and permeabilize the RNA-encasing membrane. This molecular beacon (MB) mixture was further diluted 25-fold into the purified biofluid sample and incubated at 37 °C in the dark for 2 hours to promote hybridization of the beacons to the target miRNA.

### Capture of extracellular vesicles (EVs) and lipoprotein particles

A 3 % (w/v) solution of BSA in PBS was incubated in each well at room temperature for 1 hr to block non-specific particle capture. After the removal of BSA, the biofluid samples containing EVs and lipoproteins (including pre-hybridized and untreated samples) were subsequently incubated in the wells for 2 hr at room temperature in a dark environment. For the untreated samples, excess EVs and lipoproteins were washed by pipetting PBS up and down 10 times for a total of 3 repetitions. For the pre-hybridized samples, PBS was incubated in the wells for 5 min then pipetted up and down 10 times to remove excess EVs and lipoproteins and unhybridized molecular beacons. The rinsing step was repeated 4 times in a dark environment.

### Immunofluorescence of protein markers

To prevent non-specific adhesion of the fluorescent-dye-conjugated antibodies, each well was first incubated with a 3% (w/v) BSA solution in PBS at room temperature for 1 hour. After removing the BSA solution, a 1 µg/mL solution of fluorescent-dye-conjugated antibodies (Table 2) in 10% (w/v) normal goat serum (NGS; Thermo Fisher Scientific) was added to each well and incubated for 1 hour at room temperature in the dark. Excess fluorescent antibodies were removed by washing with PBS for 3 times.

### Qualitative and quantitative analysis of the biochip

A 10 × 10 array of images was captured for each well using total internal reflection fluorescence microscopy (TIRFM; Nikon, Melville, NY) with a 100× objective and immersion oil to minimize surface refraction. Exposure times and laser power were kept constant across all experiments to maintain assay consistency. Quantification of the images involved measuring the total and mean fluorescence intensity of each spot detected by TIRFM. Histograms were created to represent the distribution of total and mean intensities of individual spots, while scatter plots displayed the relationship between mean intensity and spot size. Total fluorescence intensities of the samples were calculated using custom algorithms described in previous reports^38,56^.

## Acknowledgments

We acknowledge all the patients and healthy volunteers participating in this study. Electron microscopy was performed at the Center for Electron Microscopy and Analysis (CEMAS) at The Ohio State University. We acknowledge support from the Shared Materials Instrumentation Facility (SMIF) at Duke University.

## Author contributions

K.T.N., J.X., C.C., and F.L. were involved in conducting experiments, collecting data, analyzing results, and drafting the manuscript. M.D.H. drafted the manuscript. X.Y.R. analyzed results. J.Y.A. and B.H.M. were responsible for acquiring clinical samples from Samsung Medical Center. C.K.N. fabricated biochip. Y.K., T.J.H., L.W., D.T.W.W., and E.R. contributed to the conceptualization, design of experimental protocols, data analysis, interpretation, and the writing and review of the manuscript. Additionally, Y.K., T.J.H., D.T.W.W., and E.R. secured funding for this project.

## Data and materials availability

All data needed to evaluate the conclusions in the paper are present in the paper and/or the Supplementary Materials. Additional data related to this paper may be requested from the authors.

