## Supporting Information for "Integrated Techniques for Extracellular Particle Separation and Single-Particle Multiparametric Characterization to Track Cancer Biomarkers from Tissue to Biofluids"

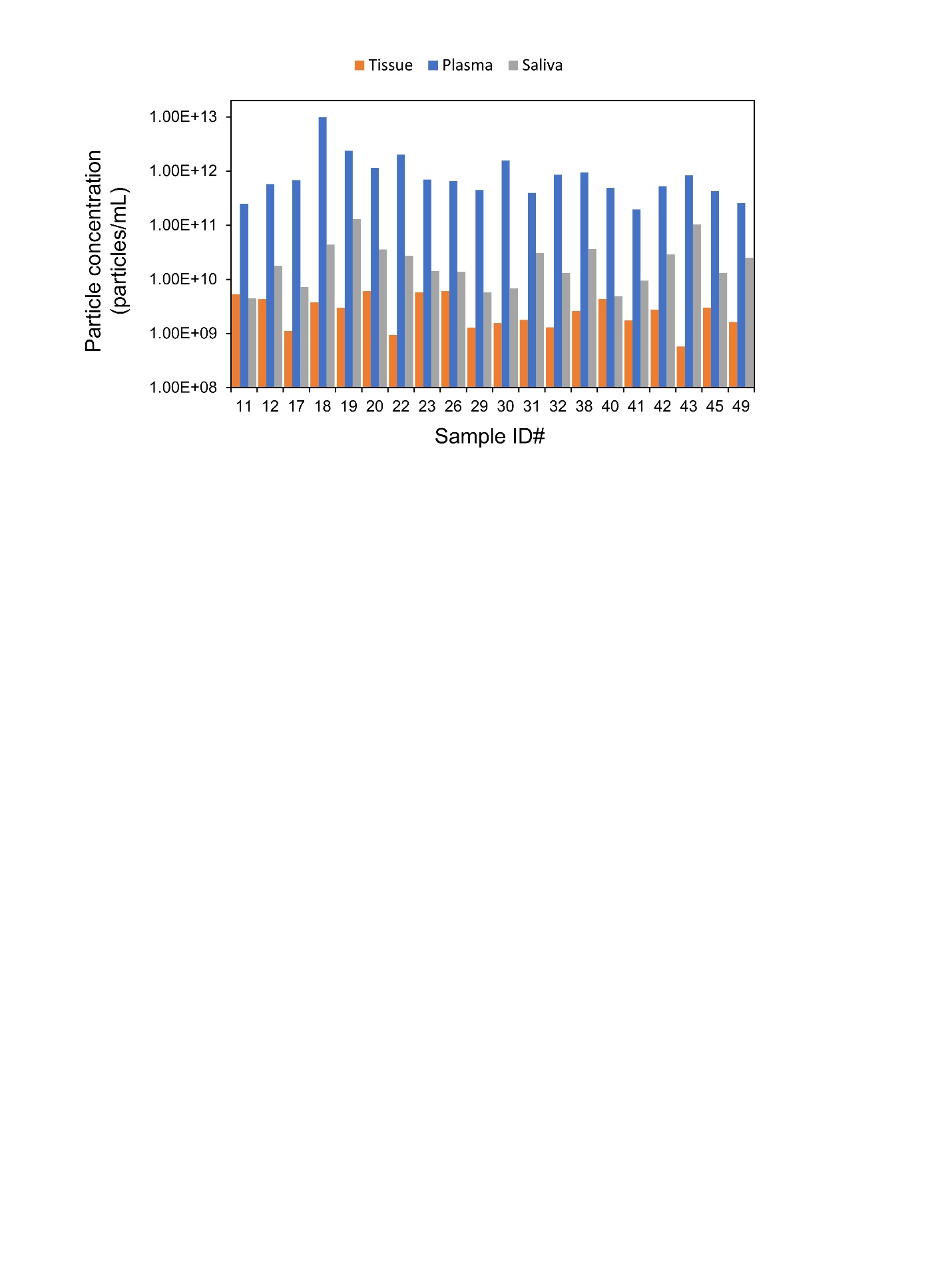

**Figure S1: Nanoparticle Tracking Analysis (NTA) measurements of extracellular vesicle particles (EVPs) isolated from plasma, saliva, and tissue samples.** The average particle concentration in plasma samples was higher compared to saliva and tissue. Prior to performing the multiplex bead assay (MBA), all samples were normalized based on their particle concentration.

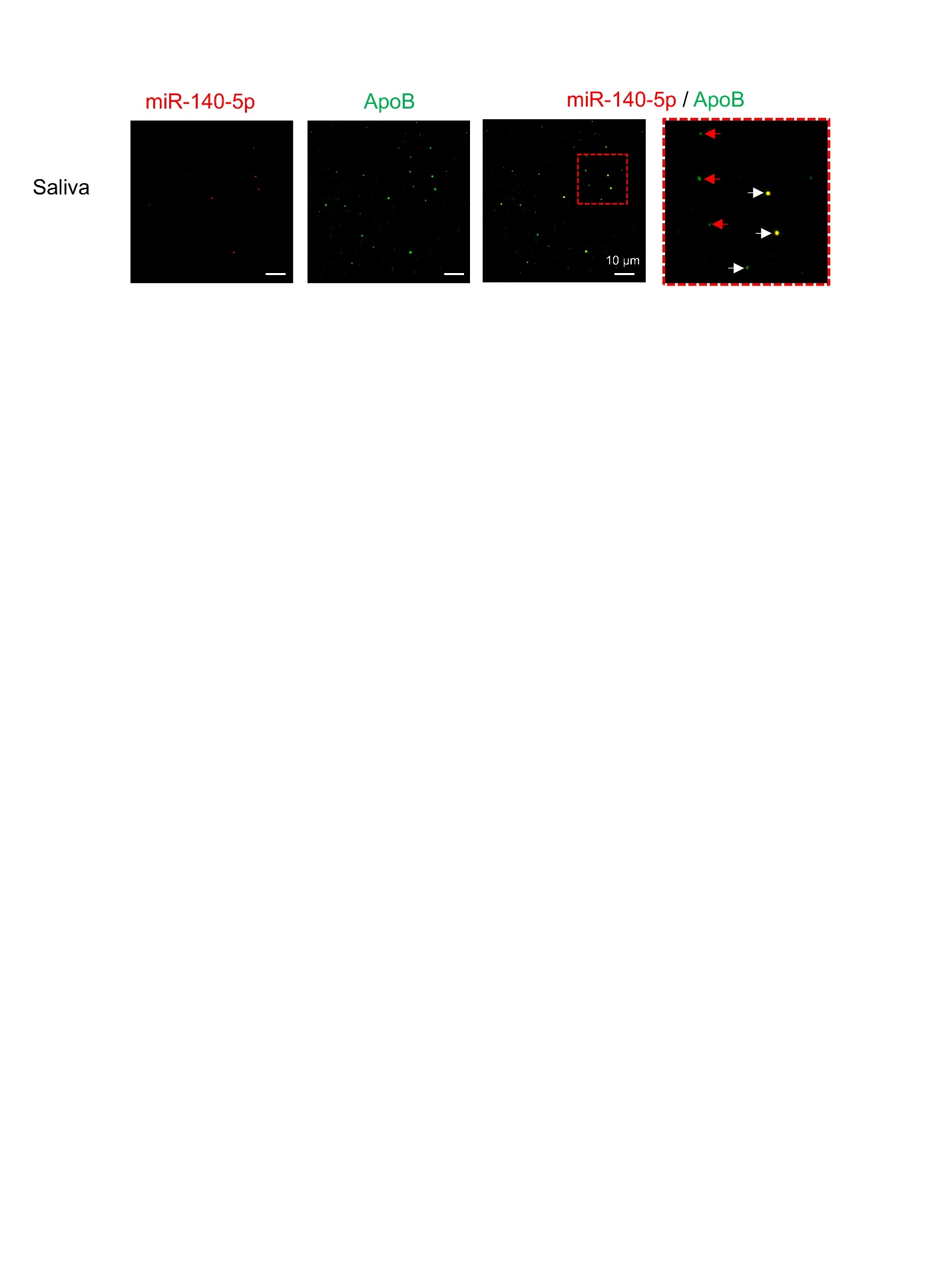

**Figure S2.** **Single lipoprotein particle multi-parametric analysis on biochip**. Representative total internal reflection fluorescence microscopy (TIRFM) images illustrating the Apolipoprotein B (ApoB) common marker of lipoprotein on single particles and miR-140-5p inside the particles. The zoom-in merged images from TIRFM illustrate the presence of anti-ApoB antibody and miR-140-5p on a single lipoprotein particle. The combined detection methods reveal colocalization of fluorescent signals within a specific localized region.

**Table S1**. **Demographic characteristics of patient and control donors enrolled in the study.**

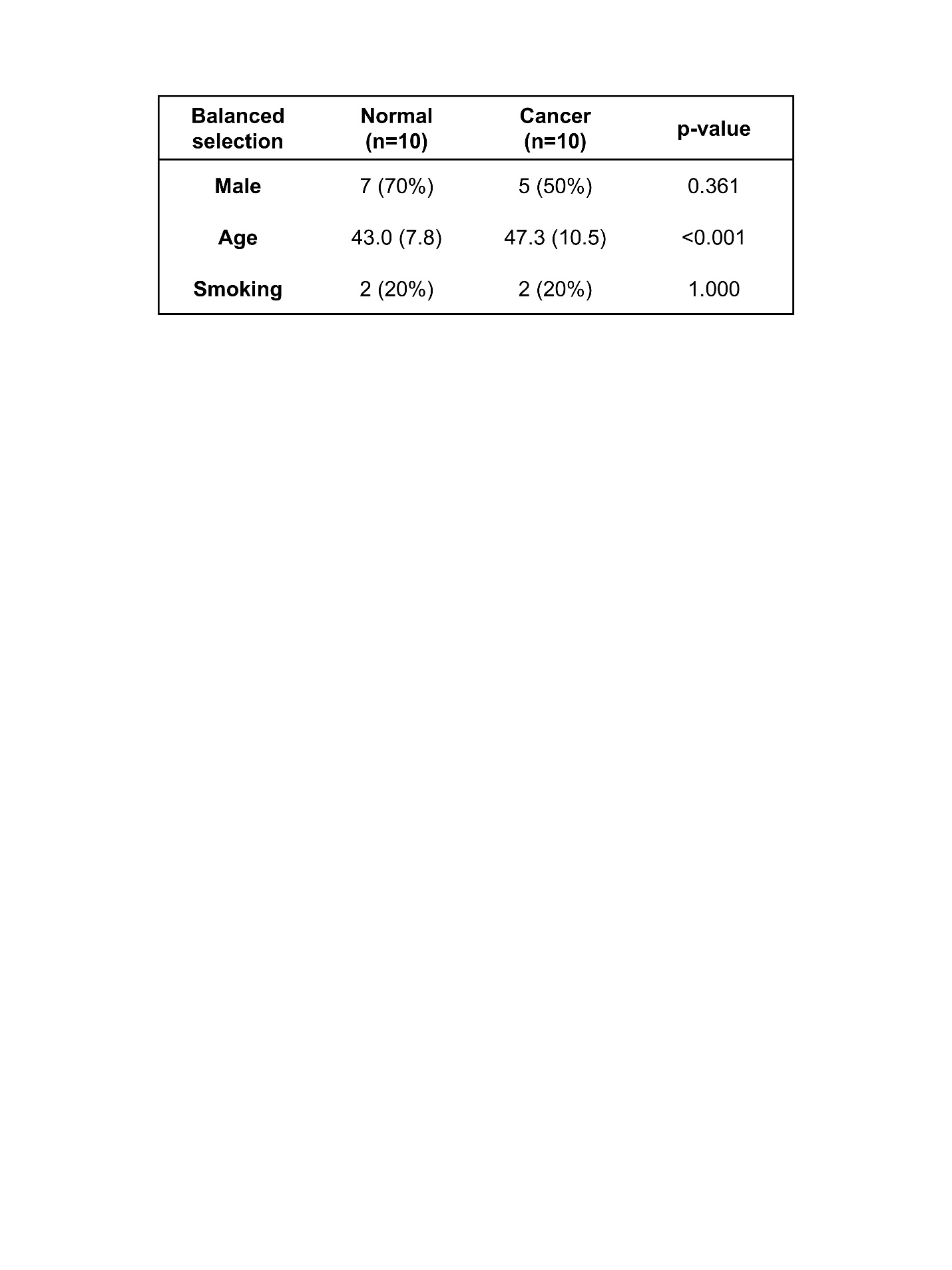

**Table S2. List of antibodies used for EVP capture and detection.**

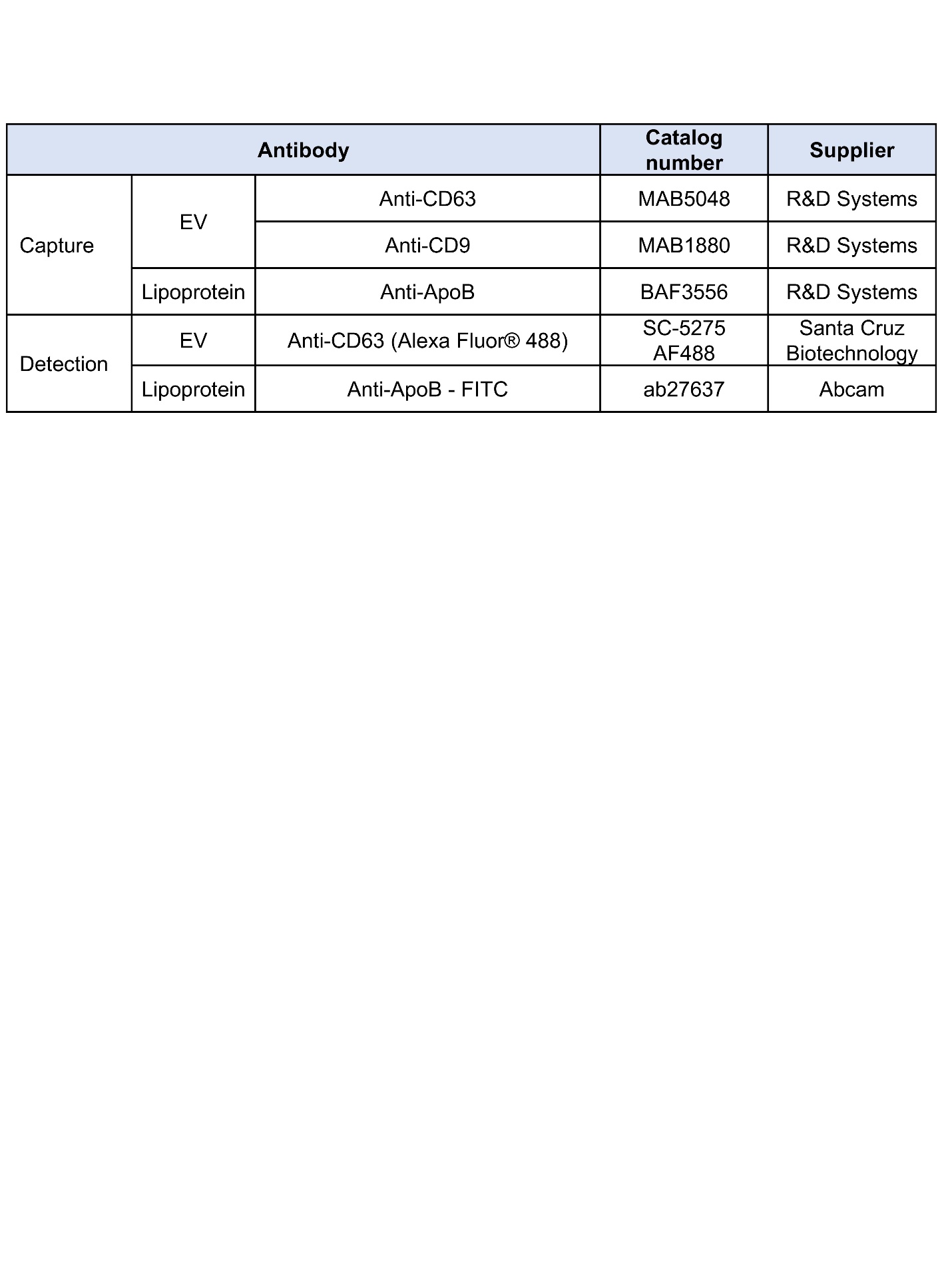

**Table S3.** **List of MB designs.**

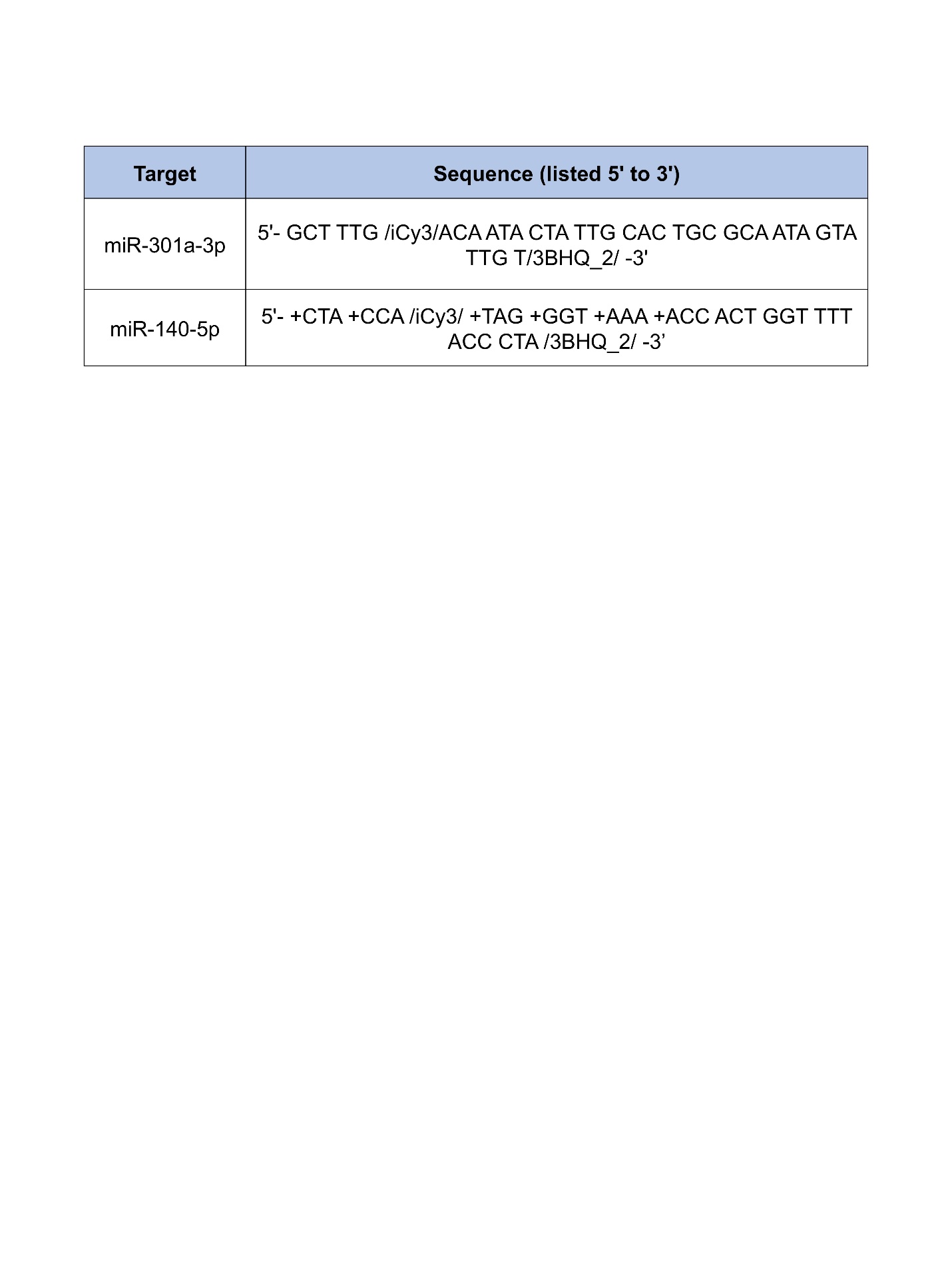
